# qtQDA: quantile transformed quadratic discriminant analysis for high-dimensional RNA-seq data

**DOI:** 10.1101/751370

**Authors:** Necla Koçhan, Gözde Y. Tütüncü, Gordon K. Smyth, Luke C. Gandolfo, Göknur Giner

## Abstract

Classification on the basis of gene expression data derived from RNA-seq promises to become an important part of modern medicine. We propose a new classification method based on a model where the data is marginally negative binomial but dependent, thereby incorporating the dependence known to be present between measurements from different genes. The method, called qtQDA, works by first performing a quantile transformation (qt) then applying Gaussian Quadratic Discriminant Analysis (QDA) using regularized covariance matrix estimates. We show that qtQDA has excellent performance when applied to real data sets and has advantages over some existing approaches. An R package implementing the method is also available.

## Introduction

Classification on the basis of gene expression data has the potential to become an important part of modern medicine, e.g. for disease diagnosis and personalization of treatment. For example, consider breast cancer. This is a heterogeneous disease consisting of several distinct types, with each type being characterized, not necessarily by its morphological or clinical characteristics, but by its molecular characteristics, thereby making it difficult to diagnose the particular type affecting a patient (Perou et al., 2000). Moreover, the most effective treatment for each of these types may differ, e.g. breast cancers that are growing in response to HER2 (human epidermal growth factor receptor 2 protein) can be treated with the targeted therapy drug trastuzumab, while ER+ (oestrogen hormone receptor positive) cancers may respond to hormone therapy that blocks oestrogen, on the other hand, triple negative (hormone receptor negative and HER2 negative) cancers do not respond to targeted therapy nor hormone therapy but respond to chemotherapy. Thus, if a woman has breast cancer, it is important to classify what type of cancer she has; this knowledge allows her treatment to be personalized, increasing her chances of survival. One promising idea for achieving such classifications is to measure the pattern of gene expression in a patient sample and use this pattern of expression as data to classify which cancer type the patient has.

There are many ways of measuring gene expression. One common approach, due to its numerous advantages, is RNA-sequencing (RNA-seq) which measures gene expression across the whole genome simultaneously (see Mardis, 2008; Wang et al., 2009). RNA-seq involves three main steps: (1) mRNA is obtained from a sample and broken into millions of short segments; (2) these mRNA segments are converted into cDNA; and (3) these cDNA segments are sequenced using next-generation sequencing. The resulting sequence data is then mapped to genomic regions of interest, typically genes, and the number mapping to each region is counted. Thus, in essence, RNA-seq data consists of *counts*: for each gene we obtain a non-negative integer count which quantifies the gene’s expression level; roughly speaking, the larger the count the higher the level of expression.

Several approaches have been proposed for classifying RNA-seq data. General machine learning approaches have been investigated, e.g. support vector machines (SVMs) and k-Nearest Neighbour (kNN) classifiers, and general regression approaches have also been applied, e.g. logistic regression (see Tan et al., 2014; Zararsiz et al., 2017). Others have focused on modelling the data more directly. For example, Witten (2011) proposed the PLDA method, which models the counts using the Poisson distribution, while Dong et al. (2016) proposed the NBLDA method, which instead models the counts using the negative binomial distribution, thereby taking into account the overdispersion known to be present in RNA-seq data on biological replicates. Others still have proposed transforming the counts, e.g. using a log transformation, so that variations on traditional classification techniques become available, e.g. Gaussian classification. The best example of this sort is the method voomDLDA (Zararsiz et al., 2017). One common feature of these direct modelling approaches is that they are, in classification terminology, “naive”: they assume that measurements on the features used for classification, i.e. the genes, are statistically *independent*.

However, this independence assumption is very unrealistic, since genes are typically involved in networks and pathways, implying that a particular gene’s expression level is likely to be correlated with the expression level of other genes. Moreover, some have argued, e.g. Zhang (2017), that the assumption of independence has a non-ignorable impact on our ability to classify: it causes bias in estimated discriminant scores, making classification inaccurate. Given this, some have focused on models for the data which incorporate dependence between genes. For example, Sun and Zhao (2015) proposed the SQDA method which models log-transformed counts with the multivariate normal distribution using regularized estimates of covariance matrices, which are assumed to be different for each class. More recently, Zhang (2017) developed a Bayesian approach where the data is modelled using a (multivariate) Gaussian copula.

In this paper we propose a new classification method for RNA-seq data based on a model where the counts are marginally negative binomial but dependent. Like previous work, we use the multivariate normal distribution for classification, where each class is assumed to have its own covariance matrix. However, our approach has two key differences: (1) instead of modelling log-transformed counts, we model quantile transformed counts; and (2) we use a novel application of a powerful regularization technique for covariance matrix estimation. We call the method qtQDA: quantile transformed (qt) Quadratic Discriminant Analysis (QDA). We demonstrate the performance of the method by applying it to several real data sets, showing that it performs better than, or on par with, existing methods. qtQDA has advantages over some existing approaches, and an R package implementing the method is available.

## Methodology

### The model

First we describe the model underpinning qtQDA. Suppose we wish to classify data into one of *K* distinct classes on the basis of *m* genes (i.e. features). Let 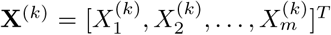 be a random vector from the *k*th class where 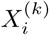 denotes the count for gene *i*. Like others, e.g. NBLDA and the method of Zhang (2017), we assume the counts are marginally negative binomial, i.e.

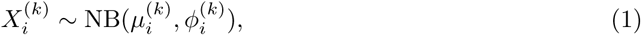

where 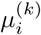 and 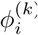 are the mean and dispersion for gene *i*, respectively (strictly speaking, 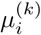 depends on the “library size”, but for the purposes of clarity, this complication is addressed later). Note that, for non-zero dispersion,

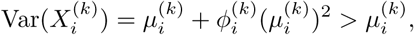

i e the data is over-dispersed relative to Poisson variation, consistent with known properties of RNA-seq data on biological replicates (see McCarthy et al., 2012). Unlike others, however, we suppose that **X**^(*k*)^ is generated by the following process:

1. Let **Z**^(*k*)^ be an *m*-vector from a multivariate normal distribution: **Z**^(*k*)^ ∼ MVN(**0, Σ**_*k*_), where *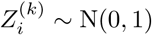*.
2. Then let the *i*th component of **X**^(*k*)^ be the transformed random variable

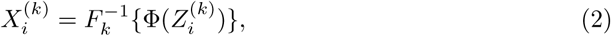

where Φ is the standard normal distribution function and *F*_*k*_ is the 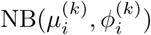 distribution function.

We make two observations. Firstly, observe that the transformation in (2) generates a vector **X**^(*k*)^ with the negative binomial margins specified in (1). This is a consequence of the following elementary fact from probability theory: if *F* and *G* are distribution functions, and *X* has distribution function *F*, then the transformed variable *G*^−1^ {*F* (*X*)} has distribution function *G* (see Lange, 2010, p. 432). We call the kind of transformation invoked here a *quantile transformation*. Note that, given the discreteness of the negative binomial distribution, the ambiguity of 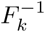 is obviated by imposing that 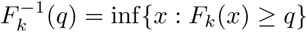, for *q* ∈ [0, 1].

Secondly, observe that the negative binomial components of **X**^(*k*)^ are not independent: the underlying MVN distribution, with a dependence structure encoded in **Σ**_*k*_, generates a dependence structure between the components of **X**^(*k*)^. Note especially that each class is assigned a different covariance matrix. As Sun and Zhao (2015) have suggested, since the presence of disease, and different disease types, leads to “rewiring” of genetic networks, and hence changes in gene associations, assuming a different covariance matrix for each class is likely to lead to better classifications. Finally, note that while the model specified by the process above is reminiscent of the Gaussian copula model of Zhang (2017), the two models are quite different.

### Classification

We now turn to how the model above is used for classification. Suppose we observe 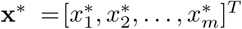 from unknown class *y*^*^, where *y*^*^ ∈ {1, 2, …, *K*}. For each class we apply the inverse of the quantile transformation (2) to the components of **x**^*^ to produce a new vector **z**^*(*k*)^, i.e. where

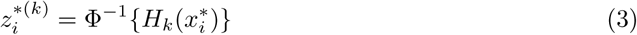

and *H*_*k*_ is a continuity-corrected version of *F*_*k*_. Here *H*_*k*_ is defined by

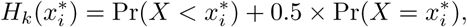

where *X* is a NB 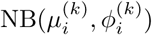 distributed random variable, and *H*_*k*_(*X*) is more nearly uniformly distributed than *F*_*k*_(*X*) itself (Routledge, 1994). The transformation from 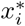 to 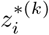 is implemented by the `zscoreNBinom` function in the R package `edgeR` (see below).

Once this transformation has been made, given the assumptions of the model, traditional quadratic discriminant analysis now becomes available, as follows. Under the model, if **x**^*^ is from the *k*th class then **z**^*(*k*)^ is an observation from the MVN(**0, Σ**_*k*_) distribution. Thus, by Bayes theorem, the posterior probability that **x**^*^ belongs to the *k*th class is

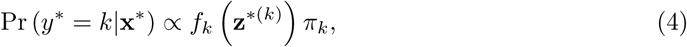

where *π*_*k*_ is the prior probability that Pr(*y*^*^ = *k*), and *f*_*k*_ is the density

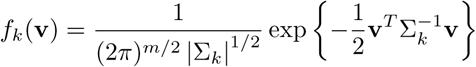

evaluated at **z**^*(*k*)^. We classify **x**^*^ into the class that maximizes this posterior probability. It is worth noting that since maximizing (4) is equivalent to maximizing log Pr (*y*^*^ = *k*|**x**^*^), this classification rule entails the following (quadratic) discriminant function:

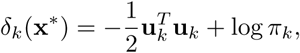

where 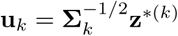, which has the following insightful interpretation: for a given class *k*, the further the vector **u**_*k*_ is from the origin, the less likely **x**^*^ is to belong to that class.

### Parameter estimation

To use the classifier in practice the parameters of the underlying model need to be estimated, i.e. the classifier needs to be “trained”. Specifically, for each gene *i* = 1, 2, …, *m* and class *k* = 1, 2, …, *K*, we need to estimate the negative binomial means 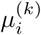 and dispersion 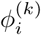, to parametrize the quantile transformation (3), and we need to estimate the covariance matrix **Σ**_*k*_ of the transformed variables, so QDA can be performed with (4). For each class *k*, suppose we have a set of *n* RNA-seq samples 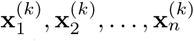 known to belong to class *k*.

To estimate the negative binomial parameters we use the methodology implemented in the R package `edgeR` (McCarthy et al., 2012; Chen et al., 2014) which is extremely fast and reliable, and offers three sophisticated approaches for dispersion estimation. Maximum likelihood estimates (MLEs) of the gene means are found by fitting a negative binomial generalized linear model (GLM) with logarithmic link function:

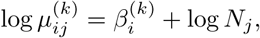

where log *N*_*j*_ is a model offset and *N*_*j*_ is the “library size” for sample *j*, i.e. the total counts 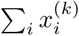 across all observed genes in the RNA-seq sample. The resulting gene mean estimates are then given by 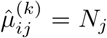 exp 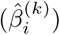. Note that the use of a GLM with log *N*_*j*_ as an offset allows us to avoid the use of “size factors” which are commonly employed in other Poisson or negative binomial based methods to scale counts to account for differences in library sizes (e.g. PLDA, NBLDA, and the method of Zhang, 2017).

The dispersion 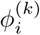 for each gene is estimated using the Cox-Reid adjusted profile likelihood (APL) function:

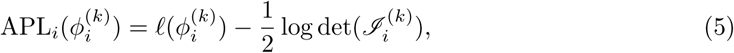

where *ℓ* is the log-likelihood and 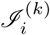 is the Fisher information of 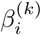, both functions being evaluated at the MLE 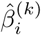. This modified likelihood function adjusts for the fact that the gene mean is estimated from the same data, thereby reducing the bias of the MLE of 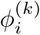. Instead of simply maximizing (5), however, to achieve even better dispersion estimates, an approximate empirical Bayes strategy is applied, where the APL for each gene is substituted by a weighted sum of APLs from carefully chosen sets of genes, resulting in “information sharing” between genes, and thereby better dispersion estimates for individual genes (see Chen et al., 2014 for details). Using different variations of this general approach, `edgeR` offers three kinds of dispersion estimates: “common”, “trended”, and “tag-wise”. By default, qtQDA uses the “tag-wise” dispersion estimates (but, the user is free to choose any of these kinds).

Once the negative binomial parameters have been estimated we apply the quantile transformation (3) to the components of the RNA-seq sample vectors to produce a corresponding set of transformed vectors 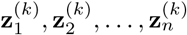 where, under the assumed model, 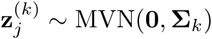. To estimate the covariance matrix **Σ**_*k*_, we begin by calculating the standard estimate:

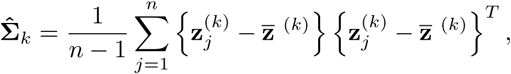

where 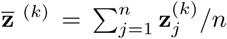. As it stands, however, this estimate is not useful in the present context where the data is typically “high dimensional”, i.e. where the number of genes used for classification will be approximately the same or greater than the number of samples (i.e. *m* ≈ *n* or *m* > *n*). In such situations this standard covariance matrix estimate is known to perform poorly (see Tong et al., 2014). To remedy this, we regularize the standard estimate using the approach developed in Schäfer and Strimmer (2005) and Opgen-Rhein and Strimmer (2007) which is implemented in the R package `corpcor` (see also Strimmer, 2008). The `corpcor` method separately shrinks the corresponding correlation estimates 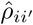 toward zero and the variance estimates 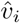 toward their median to produce the regularized estimates

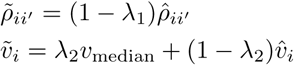

where the shrinkage intensities are estimated via

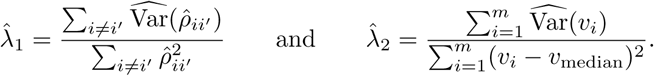

The regularized covariance matrix estimate 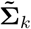 then has entries 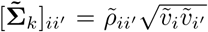. This estimate has two excellent statistical properties: (1) it is always positive definite and well conditioned (making the inverse computable); and (2) it is guaranteed to have minimum mean squared error, which is a consequence of an important result proved by Ledoit and Wolf (2003). Moreover, since the shrinkage intensities are calculated with analytic formulas, the estimate also has two significant practical advantages: (1) it is computationally very fast to compute; and (2) it does not require any “tuning” parameters. We note that the `corpcor` approach to covariance matrix regularization is quite different to the computationally intensive approach used in SQDA (Sun and Zhao, 2015). Note also that, while the `corpcor` method has previously been used for classification of gene expression data from microarrays (see Xu et al., 2009), we appear to be the first to use it for RNA-seq data.

The final parameter needed for classification with Bayes theorem (4) is the prior probability of belonging to the *k*th class *π*_*k*_ = Pr(*y*^*^ = *k*). This probability can either be specified by the user, e.g. if epidemiological knowledge is available, or estimated directly from the training data using

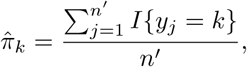

where *I* {·} is the indicator function, and *n*′ is the total number of samples in all *K* classes.

### Feature selection

Lastly, we turn to the question of which genes to use for classification. When RNA-seq is performed we typically obtain data on more than 20,000 genes. A vast number of these genes, however, will not be informative for the purposes to distinguishing between different classes. We therefore employ the following simple strategy for selecting *m* genes for classification: (1) we filter genes with low expression across all samples; (2) for each remaining gene we perform a likelihood ratio test (LRT) to test for genes differentially expressed between groups; (3) a list of genes is made, sorted by LRT statistic; (4) finally, the top *m* genes from this list is used for classification. As with negative binomial parameter estimation, this strategy is implemented using `edgeR`. Others have adopted essentially the same gene selection strategy, e.g. NBLDA, SQDA, and the method of Zhang HYPERLINK \l “bookmark32” (2017).

## Results

To assess the performance of qtQDA we apply it to three publicly available data sets:

1. *Cervical cancer data* (see Witten et al., 2010). This consists of two classes, cancer and non-cancer, each with 29 samples. Each sample consists of counts for 714 different microRNAs obtained using RNA-seq.
2. *Prostate cancer data* (see Kannan et al., 2011). This consists of two classes, 20 samples from cancer patients and 10 samples from benign matched controls. Each sample consists of RNA-seq data for the whole transcriptome.
3. *HapMap data* (see Montgomery et al., 2010; Pickrell et al., 2010). The data considered here consists of two of the HapMap populations: CEU (Utah residents with Northern and Western European ancestry) and YRI (Yoruba in Ibadan, Nigeria). There are 60 CEU samples and 69 YRI samples, each consisting of RNA-seq data for the whole transcriptome, and all from “healthy” individuals.

These data sets are very common in the RNA-seq classification literature (e.g. see Witten, 2011; Tan et al., 2014; Dong et al., 2016; and Zhang, 2017). Using these data sets, we also compare the performance of qtQDA to a number of general machine learning classifiers and specialized RNA-seq classifiers (corresponding R packages used for our analysis are listed in brackets):

- SVM (`e1071`)
- kNN (`e1071`)
- Logistic regression (`glmnet`)
- PLDA (`PoiClaClu`)
- NBLDA (http://www.comp.hkbu.edu.hk/xwan/NBLDA.R)
- voomDLDA (`MLSeq`)
- SQDA (`SQDA`)

For logistic regression, we use the GLMnet method proposed in Friedman et al. (2010) since this is one of the best representatives of this approach. This method uses a “lasso” (i.e. *ℓ*_1_) penalty in the log-likelihood function which thus overcomes many of the problems with logistic regression in high-dimensional settings (see Tan et al., 2014) and encourages regularized regression coefficients, i.e. shrunken to zero. For the SVM method we used a radial basis kernel, and for the kNN method we used k = 1, 3, and 5 (but only report results for k = 1 since this consistently performed best), and both methods were applied to log transformed counts. We apply all methods as recommended in their documentation and any “tuning” parameters were chosen with the cross-validation tools provided in the corresponding software package or chosen with our own cross-validation. The Gaussian copula method of Zhang (2017) has no publicly available implementation.

For evaluation, we estimated the true error rate, i.e. the rate at which false classifications are made, using the following bootstrap procedure: (1) each data set is randomly divided into two parts, one part consisting of 70% of the data, put aside for training the classifier, and one part consisting of 30% of the data, used as a test set to apply the trained classifier from which an error rate is recorded; (2) this is repeated 1,000 times and the error rates from each iteration is averaged to produce an estimate of the true error rate. This is the same procedure used by Dong et al. (2016) and Zhang (2017). We estimated the error rates for *m* = 100, 200, 300, 500, 700 genes, where these genes are selected using the procedure detailed in the previous section.

Results are shown in Figure 1 and Table 1. We see that qtQDA performs best for both cancer data sets, achieving the lowest error rate at 200 genes for the cervical cancer data and 100 genes for the prostate cancer data. Interestingly, for the cervical cancer data, qtQDA uniformly achieves the smallest error rate. For the HapMap data, qtQDA essentially performs as well as the SVM, kNN, and logistic regression classifiers. We note that even though these classifiers have similar performance, we think qtQDA or logistic regression would be preferred, at least in a medical context, since these classifiers do more than merely assign a sample to a particular class: they also provide a posterior probability of belonging to each class. This is important in a medical context where the different treatments or further diagnostic procedures which could be prescribed, following a classification, may be associated with very different risks.

**Figure 1.**
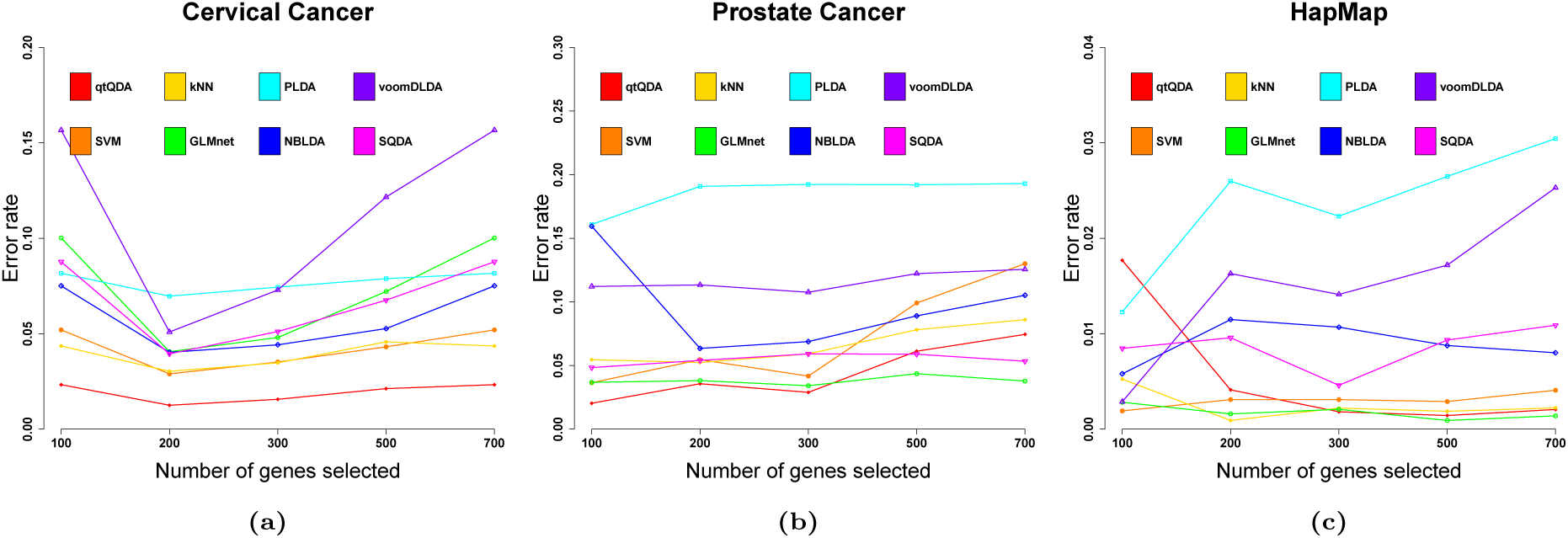
Error rate vs genes selected. These plots show classification error rate as a function of the number of genes chosen for classification for the (a) cervical cancer, (b) prostate cancer, and (c) HapMap data sets.

**Table 1.**
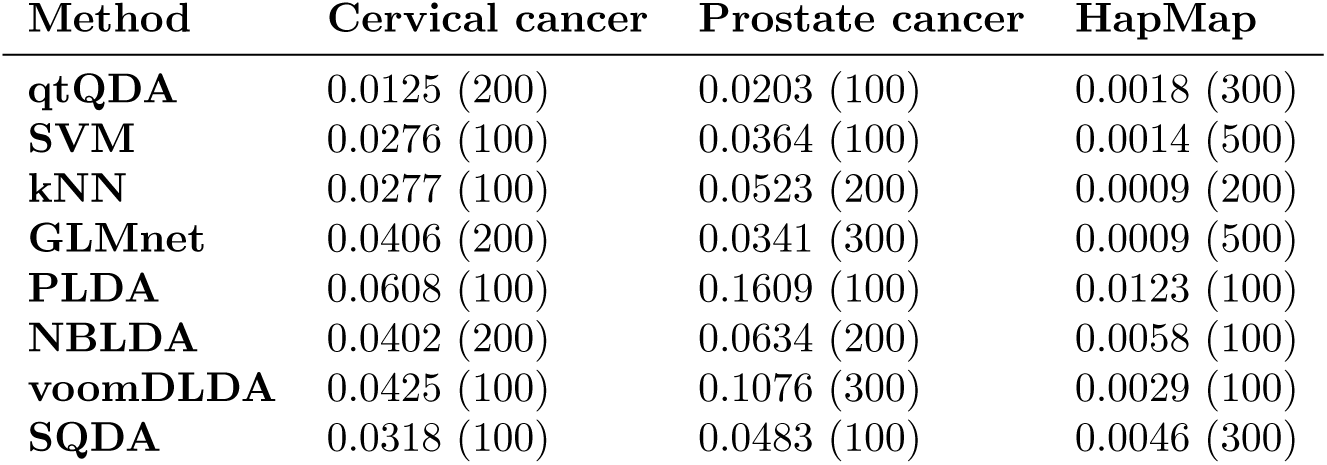
Minimum error rates. This table shows the minimum error rates achieved for each classifier in each data set. The number of genes used to obtain this minimum error rate is reported in brackets.

## Discussion

Early investigations into classification with gene expression data from microarrays, e.g. Dudoit et al. (2002), showed that making the (unrealistic) assumption of independence between measurements from different genes can still lead to classifiers with good performance. Our results, however, seem to suggest that incorporating dependence between genes can lead to even better performance, at least for RNA-seq data.

Our method has two key advantages. Firstly, unlike some approaches (e.g. kNN, GLMnet, PLDA, SQDA), qtQDA does not have any “tuning” parameters which need to be chosen with cross-validation, thus making it more straightforward to apply in practice. Secondly, in comparison to approaches which take gene dependence into account, e.g. SQDA and the method of Zhang (2017), qtQDA is computationally much faster. SQDA adopts a computationally intensive method for covariance matrix regularization. In an effort to reduce the required computation, the authors impose a block diagonal structure on the covariance matrix where each block is assumed to be the same size (but which needs to be determined by cross-validation), simplifications which even the authors acknowledge are unrealistic (e.g. under these assumptions the order of the genes used for classification matters). Yet, despite these simplifications, extensive computation is still required, making the method very slow. On the other hand, the regularization approach applied in qtQDA requires no special assumptions for the covariance matrix and requires minimal computation since the regularized estimate is obtained with analytic formulas. The Gaussian copula method of Zhang (2017) is also computationally intensive, but for a different reason: it is cast in a Bayesian framework and requires a Metropolis-Hasting algorithm, in combination with Gibbs sampling, for parameter estimation. As the author acknowledges, the computations required are time consuming even when implemented in a fast language like C++.

As Dudoit et al. (2002) points out, there are three related statistical problems in the area of classifying disease with gene expression data: (1) identifying new disease subclasses, i.e. cluster analysis; (2) classifying samples into known disease classes, i.e. discriminant analysis; and (3) identifying “marker” genes that characterize different disease subclasses, i.e. variable selection. This paper has firmly focused on problem (2), which is why it was sufficient to evaluate classifier performance solely in terms of error rate and not sparsity, i.e. the number of features used to make classifications. The feature selection method we proposed, while likely to deliver many genes informative for classification, is clearly too simplistic to deliver *only* those genes which are informative for distinguishing between classes. Thus, future research will aim at developing a sparse version of qtQDA, involving some level of regularization for features, i.e. identifying less informative features and reducing their influence to zero (e.g. like the GLMnet logistic regression classifier). A sparse qtQDA may also help address problem (3) above, the answer to which has practical advantages, e.g. knowing which subset of genes need to be measured for effective classification, and theoretical advantages, e.g. obtaining insight into the underlying biological process driving the disease (or subclass) in question. A sparse qtQDA may also deliver a further bonus: it may lead to a better answer to problem (2), i.e. to even better disease classifications.

## Conclusion

We have proposed a new classification method for RNA-seq data based on a model where the data is marginally negative binomial but dependent, thereby incorporating dependence between genes. The method works by first performing a quantile transformation then applying Gaussian quadratic discriminant analysis, where each class is assumed to have its own covariance matrix. The classifier is trained by using the sophisticated `edgeR` methodology for negative binomial parameter estimation, to parametrize the quantile transformation, and by using the powerful `corpcor` methodology for regularized covariance matrix estimation, so that effective quadratic discriminant analysis can be performed on the transformed data. We have shown that, when applied to real data sets, the classifier has excellent performance in comparison to other methods, and has two key advantages which makes it easy to apply in practice: (1) it does not have any tuning parameters; and (2) it is computationally very fast. An R package implementing the method is also available.

## Appendix

### Software availability

An R software package implementing the qtQDA method is available via the following link: https://github.com/goknurginer/qtQDA

### Author contributions

The project was conceived by GG, GYT, and NK. The method was proposed by LCG and GKS, with contributions from NK. All data analysis was performed by NK, under the supervision of GG, LCG, and GYT. The software implementation was designed and written by NK, GG, and GKS, with contributions from LCG. The manuscript was written by NK and and LCG, with contributions from GKS.

## Acknowledgements

We thank Prof. Terry Speed for helping us clarify the differences between our qtQDA model and the Gaussian copula model of Zhang (2017), for recommending the `corpcor` covariance matrix regularization method, and for commenting on a draft manuscript.

## Additional information and declarations

This work was supported by The Scientific and Technical Research Council of Turkey (TUBITAK2214/A - 1059B141601270 to NK), and The Australian National Health and Medical Research Council (Program Grant 1054618 and Fellowship 1154970 to GKS), the Cancer Therapeutics CRC, Victorian State Government Operational Infrastructure Support and Australian Government NHMRC IRIIS. Funding for open access charge: Smyth Lab funds.

## Notes

#### Summary of Updates

The section "Additional information and declarations" is added. Author contributions are updated. A couple of typos are corrected.

https://github.com/goknurginer/qtQDA

